# Dose escalation pre-clinical trial of novel DOK7-AAV in mouse model of DOK7 congenital myasthenia

**DOI:** 10.1101/2024.02.09.579626

**Authors:** Judith Cossins, Imre Kozma, Claudia Canzonetta, Al Hawkins, David Beeson, Patricio Sepulveda, Yin Dong

## Abstract

Congenital myasthenic syndromes (CMS) are a group of inherited disorders characterised by defective neuromuscular transmission and fatigable muscle weakness. Mutations in *DOK7*, a gene encoding a post-synaptic protein crucial in the formation and stabilisation of the neuromuscular junction (NMJ), rank among the leading three prevalent causes of CMS in diverse populations globally. The majority of DOK7 CMS patients experience varying degrees of disability despite receiving optimised treatment, necessitating the development of improved therapeutic approaches. Here we executed a dose escalation pre-clinical trial using a DOK7-CMS mouse model to assess the efficacy of Amp-101, an innovative AAV gene replacement therapy. Amp-101 is based on AAVrh74 and contains human DOK7 cDNA under the control of a muscle-restricted promoter. We show that at doses 6x10^13^vg/kg and 1x10^14^ vg/kg, Amp-101 generated enlarged NMJs and rescued the very severe phenotype of the model. Treated mice became at least as strong as WT littermates and the diaphragm and tibialis anterior muscles displayed robust expression of DOK7. This data suggests that Amp-101 is a promising candidate to move forward to clinic trials.

## Introduction

Congenital myasthenic syndromes are a group of rare inherited disorders that are characterised by fatiguable muscle weakness, which is caused by defective signalling at the neuromuscular junction (NMJ). Causative mutations in over 30 genes have been identified (Ohno et al, 2023; Rodriguez Cruz et al, 2018) including *DOK7* which encodes the downstream of kinase 7 protein (DOK7) (Beeson et al, 2006). This protein is crucial for the dimerization and activation of muscle specific kinase (MuSK), which is a tyrosine kinase that orchestrates nicotinic acetylcholine receptor (nAChR) clustering and the formation of the neuromuscular junction (Inoue et al, 2009; Okada et al, 2006). Patients with mutations in *DOK7* usually present during childhood with a familial limb-girdle myasthenia, and have characteristic small NMJs with a reduced number of post-synaptic folds (Beeson *et al*., 2006(Slater et al, 2006)).

DOK7-CMS patients do not respond well to conventional CMS therapies such as the acetylcholinesterase inhibitor pyridostigmine, but ephedrine was found to be beneficial (Lashley et al, 2010; Palace et al, 2007; Schara et al, 2009). However, due to the limited medical access to ephedrine, the preferred treatment is currently salbutamol (albuterol), which is as efficacious as ephedrine and has a good and extensive safety profile in children (Burke et al, 2013; Liewluck et al, 2011; Lorenzoni et al, 2013; Mahjneh et al, 2013). Both of these medicines are β2-adrenergic receptor agonists, and although the precise therapeutic mechanism is unclear, evidence suggests that salbutamol increases both nAChR clustering *in vitro* (Clausen et al, 2018) and, importantly, increases the post-synaptic area and post-synaptic folding *in vivo* (Vanhaesebrouck et al, 2019). NMJ size is thought to be important for efficient signalling and an increase should improve muscle strength. However, many patients show a poor response (Liewluck *et al*., 2011) and remain disabled even on optimised treatment. In addition, salbutamol causes tachycardia and muscle cramps in many patients (Lee et al, 2018), limiting the dose that can be used. Therefore, a more efficacious treatment with fewer side effects is needed.

An alternative potential treatment is gene replacement therapy whereby a functional wild type *DOK7* gene would be introduced via a recombinant adeno-associated virus (rAAV). Arimura et al. highlighted the potential of this therapy when his group successfully treated a DOK7-CMS mouse model with a rAAV9 expressing human DOK7 tagged with EGFP (Arimura et al, 2014). This knock-in mouse model is homozygous for a frameshift mutation, c.1124-1127 dupTGCC, which corresponds with the most commonly observed mutation in CMS patients. These mice have a severe phenotype, gain very little weight and only survive a few days after birth. The introduced gene generated hugely enlarged NMJs and, remarkably, the mice not only thrived but also survived for a year during the follow-up period.

rAAV expressing DOK7 has also been shown to generate enlarged synapses and ameliorate symptoms in other neuromuscular diseases. In all cases DOK7 expression was under control of ubiquitous promoters. The CMV promoter was used in models for Emery-Dreifuss muscular dystrophy (Arimura *et al*., 2014) and Amyotrophic lateral sclerosis (Miyoshi et al, 2017), and the chicken beta-actin promoter for a spinal muscular atrophy mouse model (Kaifer et al, 2020). More recently, long-term targeted expression of DOK7 in skeletal muscle of wild type mice by using the tMCK promoter also generated enlarged NMJs with no evidence of adverse pathology (Huang et al, 2023).

Using a rAAV gene transfer approach only requires a single administration, thereby circumventing the need for repeat daily dosing. However, high doses of AAV may cause liver toxicity and death, as exemplified by the death of two children with spinal muscular atrophy who received AAV9 gene therapy Zolgensma (Philippidis, 2022), one child who received AAV9 gene therapy PF-06939926 Duchenne muscular dystrophy gene therapy (Armstrong, 2021), and two patients in a recent phase 2 clinical trial of AT132, an AAV8 gene therapy in X-linked myotubular myopathy (2020). Therefore, new AAV gene therapies with a lower therapeutic dose should be preferred for a safer clinical profile.

The aim of this study was to test the efficacy of three doses of a rationally optimised DOK7-AAV construct, Amp-101, in the DOK7-CMS mouse model described above (this model will be referred to as Dok-7^KI/KI^). Amp-101 is based on AAVrh74 and contains human DOK7 under control of a muscle-restricted promoter. AAVrh74 was first found in rhesus macaques, and is potentially less susceptible to treatment blockage by pre-existing serum AAV antibodies than some of the other AAV serotypes that have been isolated from humans (Mendell et al, 2020). This, along with its ability to efficiently transduce skeletal muscle (Pozsgai et al, 2017; Sondergaard et al, 2015; Zygmunt et al, 2017), and a relatively good safety profile in a DMD clinical trial (Mendell *et al*., 2020) make it a good gene delivery vehicle for this study.

## Results

### Study plan

In this dose escalation trial we administered three different doses of Amp-101 intra-peritoneally (IP) into 4-day old Dok-7^KI/KI^ model mice: 2x10^13^ vg/kg, 6x10^13^ vg/kg and 1x10^14^ vg/kg. As controls, WT littermates were injected with saline. Table 1 shows the number of mice and group designation. The virus was diluted in sterile 0.9% w/v saline by a third party so that the experimenter was blinded to the dose until after data collection and analysis. Mice were culled at either 1 month or 3 months of age. Mouse weight and survival was monitored. For the 3-month groups, muscle fatiguability was monitored every two weeks from 30 days of age using an inverted screen hang test, which is a well-established measure of strength endurance and fatigable weakness. Strength was also assessed by rotarod and grip strength test at 2 and 3 months of age. Electromyography (EMG) was performed at 3 months of age. DOK7 expression in TA, diaphragm and triceps brachii was analysed by western blotting for the 3-month old mice, and the morphology of NMJs in diaphragm was analysed in both the 1- and 3-month groups. The primary efficacy measure was survival and the secondary outcome measure strength as measured by the inverted screen hang test. Fig. 1a shows a schematic diagram of the timeline.

**Figure 1.**
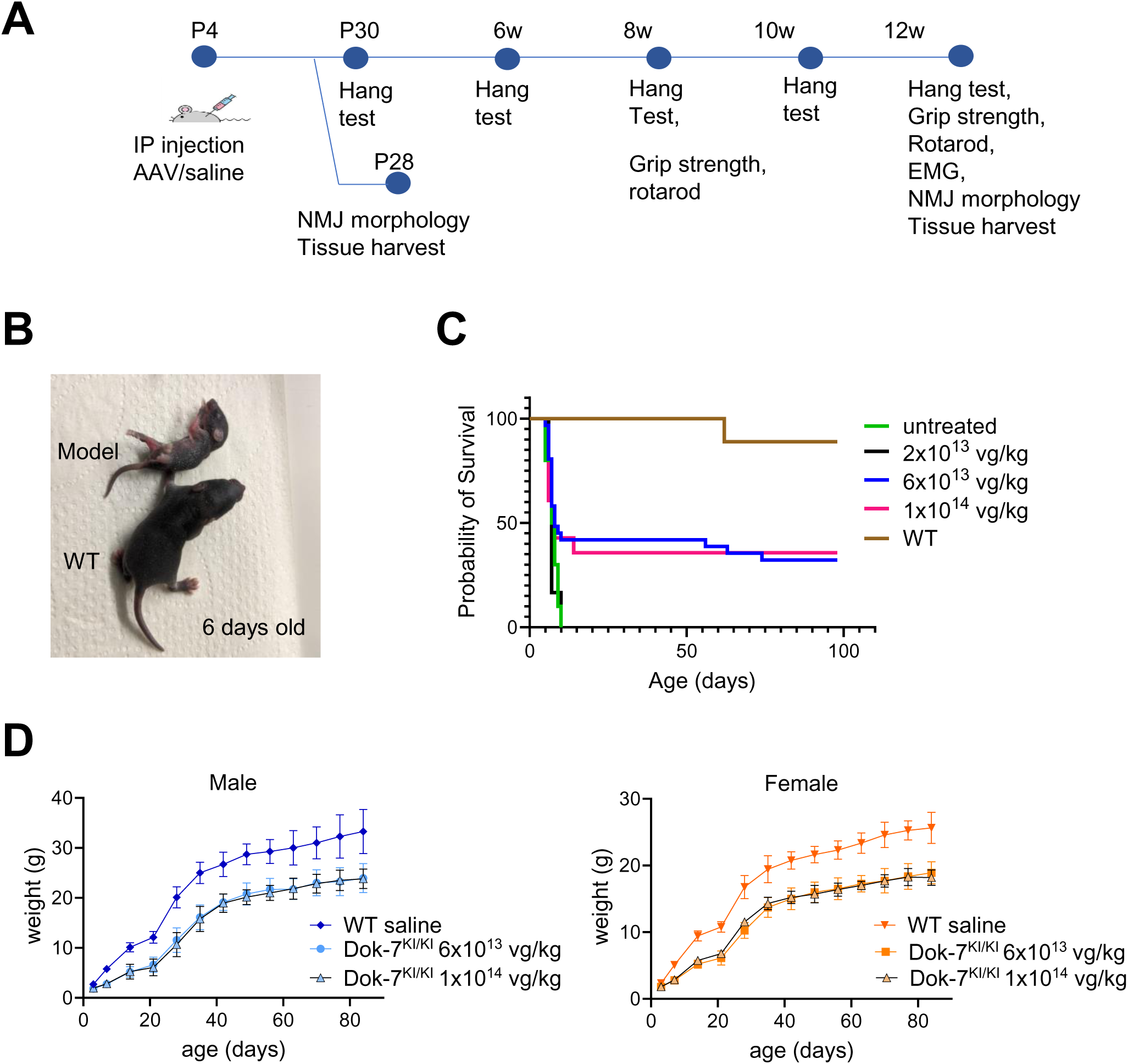
Timeline, survival and growth of Amp-101 treated Dok-7^KI/KI^ mice. A Timeline outlining the study plan, indicating when behavioural tests were carried out. B Photograph showing the size difference of an untreated Dok-7^KI/KI^ mouse and a WT littermate at 6 days of age. C Kaplan-Meier survival curve of Dok-7^KI/KI^ either untreated or treated with various doses of Amp-101 as indicated. Survival curve of saline injected WT littermates is also shown. D Male and female growth curves of Dok-7^KI/KI^ mice treated with 6x10^13^ vg/kg or 1x10^14^ vg/kg and saline injected WT littermates up to 3 months of age.

**Table 1.** Group designation and number of mice used in the study.

| Group | Test article | Dose vg/kg | Timepoints | Number of male and female mice reaching endpoint |
| --- | --- | --- | --- | --- |
| 1 | Dok-7<sup>KI/KI</sup> | 0 | survival endpoint | 5 male/5 female |
| 2 | WT age matched | Vehicle | 1 month, 3 months | 4 male/4 female, 4 male/4 female |
| 3 | Dok-7<sup>KI/KI</sup> mice | 6x10<sup>13</sup> | 3 months | 5 male/5 female |
| 4 | Dok-7<sup>KI/KI</sup> mice | 2x10<sup>13</sup> | 3 months | 5 male/5 female |
| 5 | Dok-7<sup>KI/KI</sup> mice | 1x10<sup>14</sup> | 3 months | 5 male/5 female |
| 6 | Dok-7<sup>KI/KI</sup> mice | 6x10<sup>13</sup> | 3 months | 5 male/5 female |
| 7 | Dok-7<sup>KI/KI</sup> mice | 2x10<sup>13</sup> | 1 month | 5 male/5 female |
| 8 | Dok-7<sup>KI/KI</sup> mice | 1x10<sup>14</sup> | 1 month | 5 male/5 female |

### Amp-101 treatment improved survival and weight gain at higher doses, but not at the lowest dose

The Dok-7^KI/KI^ mouse model used in this study has the common human mutation c.1124_1127dupTGCC knocked in. It has a very severe phenotype, fails to gain weight, and dies within a few days of birth (Fig EV1). Fig. 1b illustrates a comparison between an untreated 6-day-old model mouse and a wild-type (WT) littermate. The lowest dose of 2x10^13^ vg/kg did not rescue the severe phenotype of this model in the first 6 mice injected, and so this group was discontinued for ethical reasons (Fig. 1c). For the other two doses, approximately 32% of mice treated with 6x10^13^ vg/kg and 35% of mice treated with 1x10^14^ vg/kg survived beyond 10 days. After reaching 10 days of age, three male mice treated with 6x10^13^ vg/kg, one male mouse treated with 1x10^14^ vg/kg and one WT male mouse reached the humane end point before the end of the trial due to >15% weight loss compared to their previous recorded maximum (Fig. 1c). Of those mice that survived, the AAV-treated model mice gained weight at a similar rate to WT littermates after weaning, but were approximately 25% (female) or 28% (male) smaller (Fig. 1d).

### Amp-101 treated Dok-7^KI/KI^ model mice were at least as strong as WT mice for the duration of the study

Three different tests were carried out to assess the muscle strength of the AAV-treated model mice, and male and female mice were analysed separately.

On fortnightly inverted screen hang tests, models treated with 6x10^13^ vg/kg or 1x10^14^ vg/kg Amp-101 could hang on for at least as long as WT mice at all time points throughout the trial, with some gender differences. Muscle strength in treated models increased until week 6, after which there was a gradual decrease in strength which was more pronounced in the male models. While there was no significant difference between male WT and models treated with 6x10^13^ vg/kg, those models treated with the highest dose of 1x10^14^ vg/kg were significantly stronger than WT littermates until week 12. In contrast, female models treated with both doses were significantly stronger then WT littermates at weeks 8 and 10, and there was no significant difference between the two doses at any timepoint. (2-way ANOVA, Sidak’s multiple comparison’s test).

We observed an inverse correlation between weight and hang time in WT mice (Fig EV2). As the treated model mice were lighter than WT mice, this might have contributed to their performance on the inverted hang test. To take this into account we normalised the data by multiplying the hang time by mouse weight (Fig 2b). Even with this normalisation factor, the model mice are still as strong as, or stronger than, WT littermates, and the gender differences noted above were still observed.

**Figure 2.**
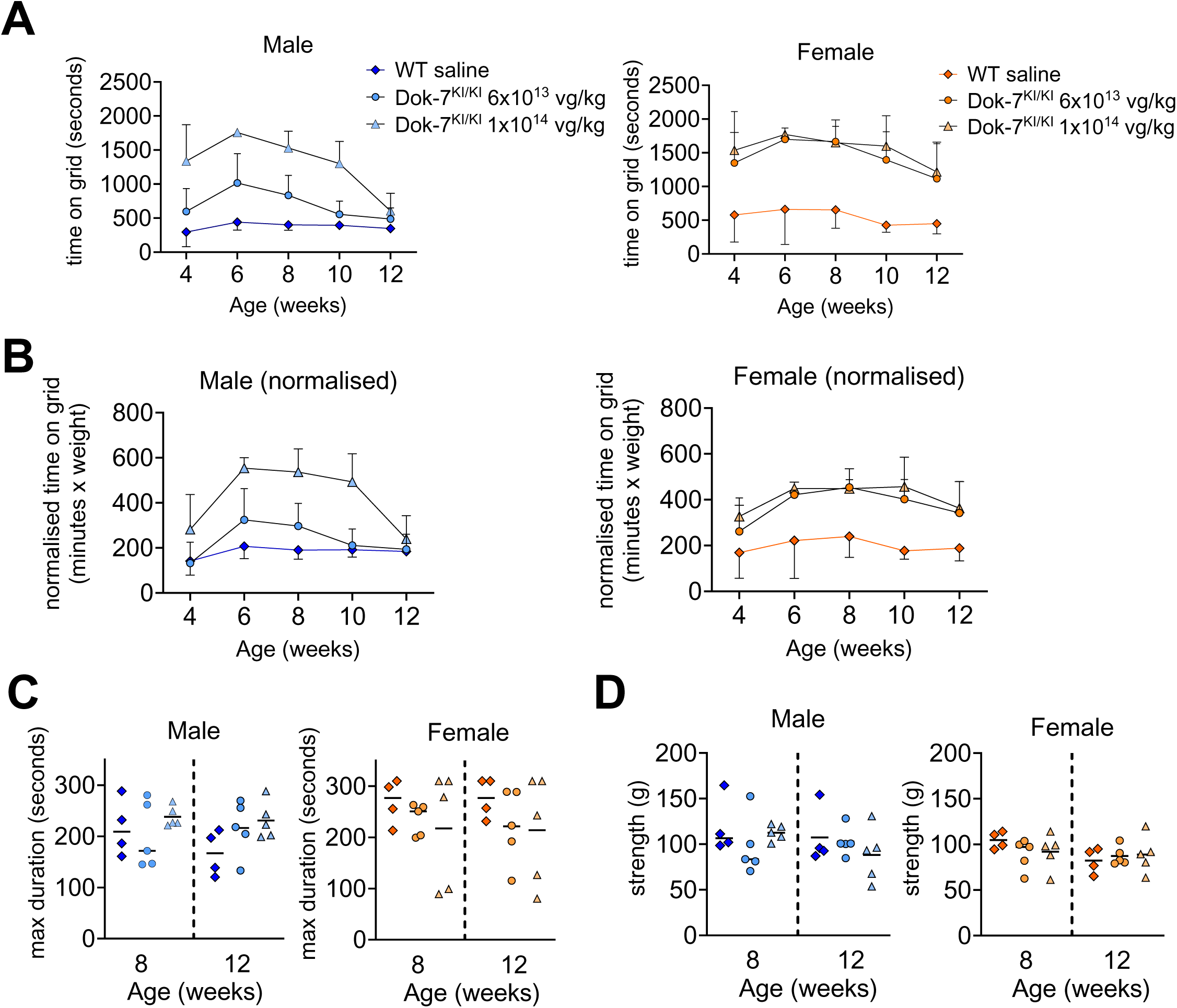
Strength tests show that Amp-101-treated Dok-7^KI/KI^ mice are as strong or stronger than WT littermates. A. Inverted screen hang test. Mice were held upside down on a grid and the time to fall was measured. Three consecutive tests were carried out for each time point and the cumulative time is shown. B. Normalised inverted screen hang test. To take into account the fact that lighter mice are able to hang onto the screen for longer, the time on grid was normalized by multiplying the time by the mouse weight. C. Rotarod test. The rotarod speed gradually accelerated from 5rpm to 40 rpm over a 5-minute period. Each test was carried out 3 times for each time point and the maximum duration is shown. D. Grip strength test. Mice were gripped by the tail and were allowed to hold onto the grid of the apparatus with their front paws. They were pulled away horizontally and the force generated by the mouse was measured. Each test was carried out 5 times for each time point and the maximum value is shown.

For rotarod and grip strength tests there was no significant difference between treated model mice and WT littermates for either dose at either age and for either gender (ordinary one-way ANOVA, Tukey multiple comparisons). In addition, within each treatment group, the strength did not significantly change between 8 and 12 weeks of age (Figs 2c and 2d show data from rotarod and grip test respectively).

### Amp-101 treated Dok-7^KI/KI^ model mice show decrement of compound muscle action potential (CMAP) on repetitive nerve stimulation

A hallmark of myasthenia is impaired neuromuscular signalling, which can be detected by electromyography (EMG). EMG involves repetitively stimulating a motor nerve (in this case the phrenic nerve) and measuring the compound muscle action potential (CMAP) after each stimulation. A decrease, or decrement, in successive CMAP amplitudes indicates defective neuromuscular signalling.

Unfortunately, due to the severity and early lethality of Dok-7^KI/KI^ mice, no EMG data could be collected before treatment, nor on untreated mice. To test whether the Amp-101 treated Dok-7^KI/KI^ mice had impaired neuromuscular signalling we carried out EMG on 3-month old mice. The sciatic nerve was given a train of 10 repetitive stimuli at different frequencies and the compound muscle action potential (CMAP) amplitude was measured at each stimulus. Each successive CMAP amplitude was then calculated as a percent of the first stimulus. Examples of this data for male WT mice and male Dok-7^KI/KI^ mice treated with 6x10^13^ vg/kg are shown in Fig 3a.

**Figure 3.**
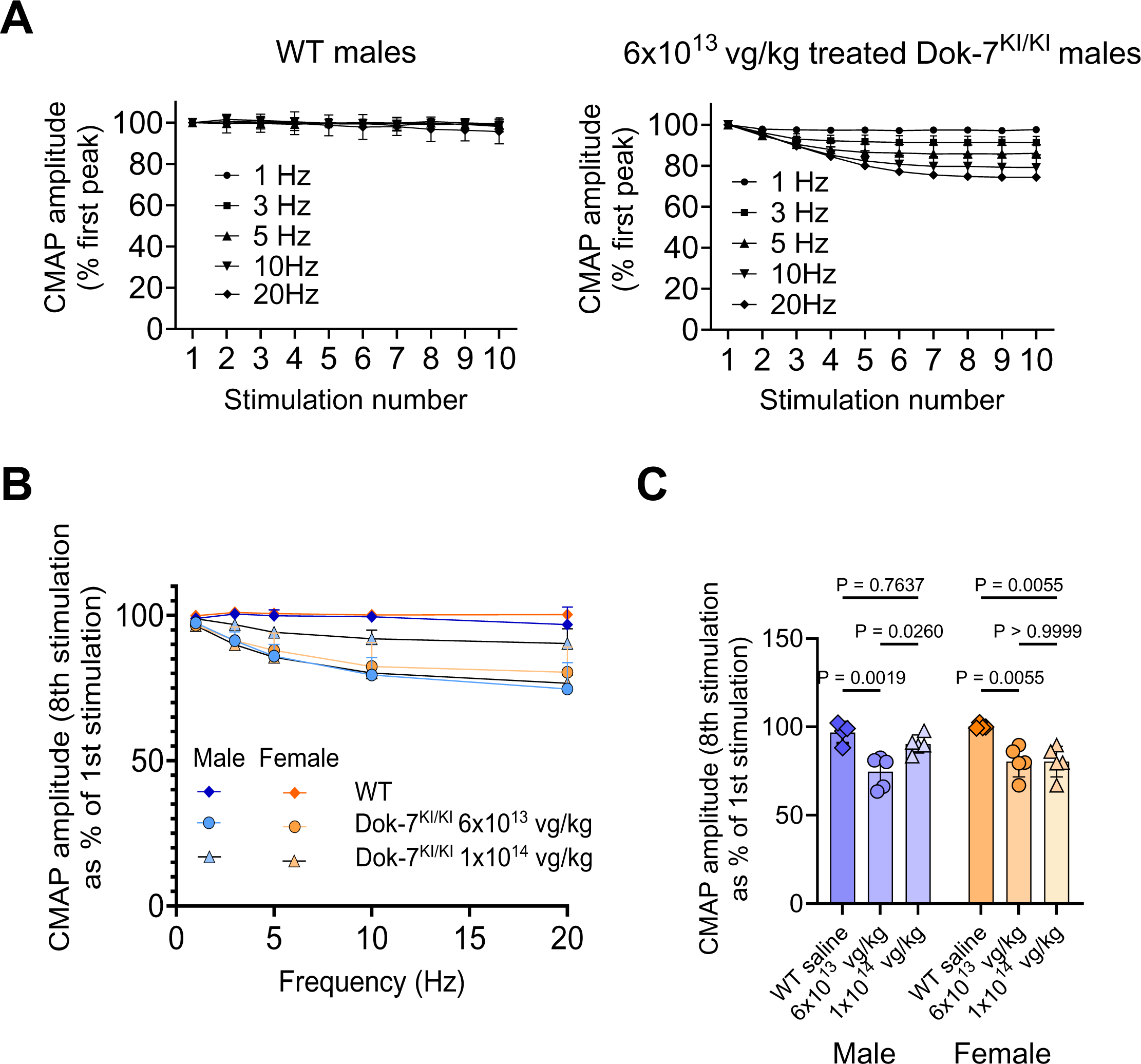
Amp-101 treated Dok-7^KI/KI^ mice show decrement of compound muscle action potential (CMAP) on repetitive nerve stimulation. A. Example traces from a WT mouse and an Amp-101 treated model mouse showing the CMAP amplitude at each stimulus in a train of ten stimuli at various frequencies as shown. B, C. (B) CMAP amplitude of the 8^th^ stimulus as a percentage of the first stimulus for each stimulation frequency is shown. The treated model mice show decrement, although the male models treated with 1x10^14^ vg/kg do not show significant decrement at 20Hz compared with WT littermates (C). P values were obtained using an ordinary one-way ANOVA with Tukey’s multiple comparisons test.

Even though the treated model mice were at least as strong as their WT littermates, there was nonetheless evidence of decrement of up to 25% at 20Hz. WT littermates did not exhibit decrement (Fig 3b). It is of note that out of the treated models, the males treated with the highest dose of 1x10^14^ vg/kg showed the least decrement (only 10% at 20Hz, Figure 3c) and this was not significantly different from WT (p=0.7637; one-way ANOVA, Tukey’s multiple comparison test).

### Enlarged NMJs are observed following treatment with Amp-101

Previous publications and our own observations have shown that overexpression of DOK7 in skeletal muscles gives rise to enlarged neuromuscular junctions (Arimura *et al*., 2014; Eguchi et al, 2020; Miyoshi *et al*., 2017; Ueta et al, 2020), and we wanted to know whether this novel rAAV construct would also give rise to enlarged synapses. Diaphragms collected from mice one or three months after treatment were stained with antibodies against neurofilament and synaptophysin to visualise the presynaptic motor neuron terminal, and post-synaptic nAChRs were stained with fluorescent α-bungarotoxin. Enlarged synapses were present in all Amp-101 treated model mice at both one and three months of age (Fig 4a shows example images). Variation in the size of NMJs on different muscle fibres in the same diaphragm was observed.

**Figure 4.**
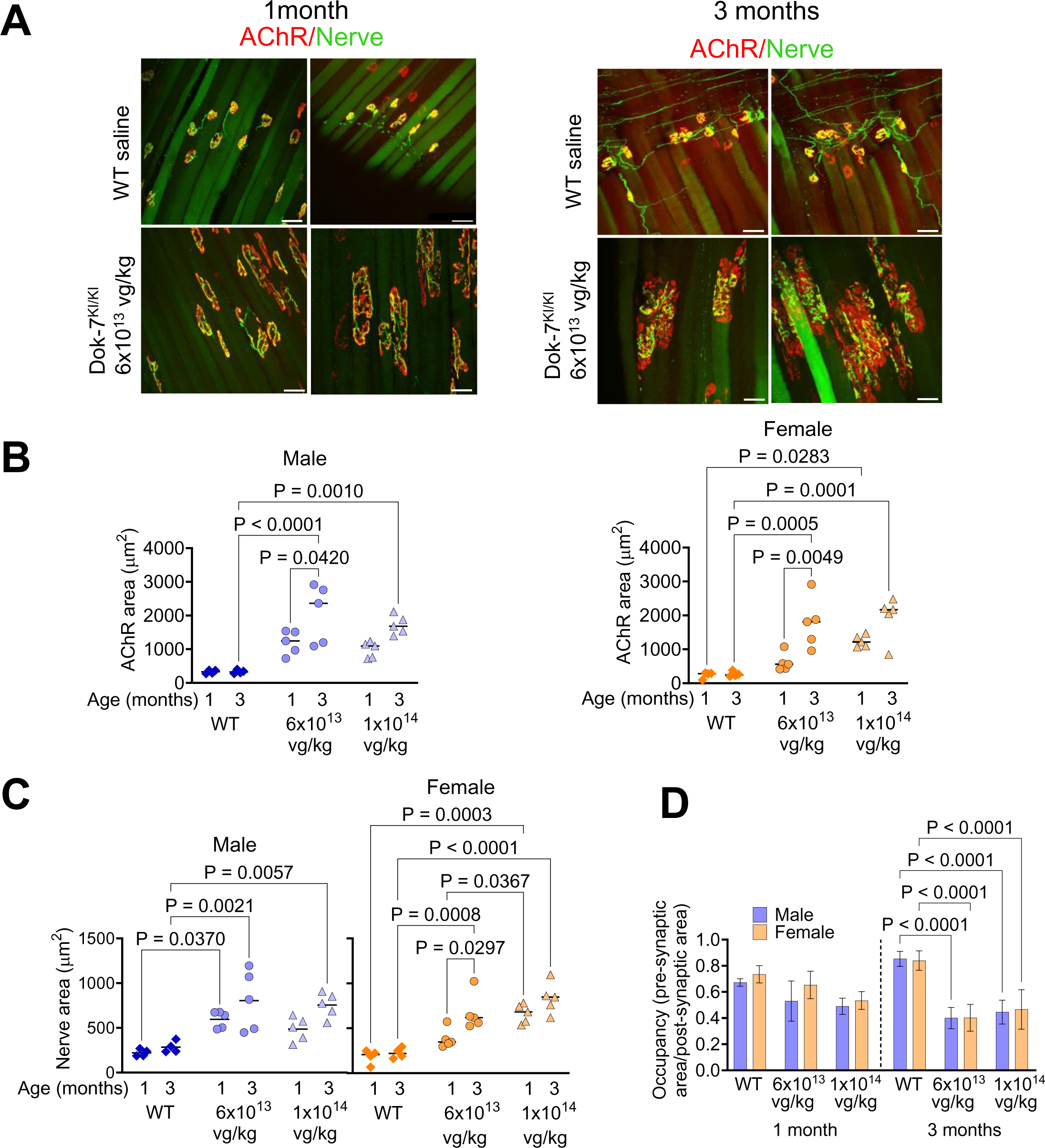
Amp-101 generates enlarged synapses in Dok-7^KI/KI^ model mice. A. Example maximum projection images generated from confocal z-stacks of diaphragm muscle from WT mice and from Dok-7^KI/KI^ mice treated with 6x10^13^ vg/kg of Amp-101. Motor neuron termini are stained with antibodies against neurofilament and synaptophysin (green) and post-synaptic nAChRs are stained with a fluorescent α-bungarotoxin (red). Scale bars are 50μm. B, C. NMJs are enlarged in Amp-101 treated model mice 1 month after treatment compared with WT littermates, and continue to enlarge as the mice grow to 3 months of age. The graphs show post-synaptic nAChR area (B) and pre-synaptic area of nerve (C), D. At 1 month of age the ratio of pre-to post-synaptic staining, known as occupancy, is not significantly different between WT and treated models. By 3 months of age occupancy is significantly lower in treated models compared with WT littermates. P values were obtained using a one-way ANOVA with Tukey’s multiple comparisons test.

Gender-specific differences were examined across various groups. In WT control mice, there was no significant disparity in the size of the NMJ between males and females at either age (Table EV1, Figure 4B). In contrast, gender-specific differences were detected in the treated model mice at 1 month of age. Specifically, in model mice treated with 6x10^13^ vg/kg, the NMJs were, on average, larger in males than females, as evidenced by both BuTx and nerve staining. However, in mice treated with 1x10^14^ vg/kg, females exhibited larger pre-synaptic areas. By 3 months of age no significant gender differences were detected in either pre-or post-synaptic staining. Given the gender disparity identified at 1 month of age, we further analysed the males and females separately.

In WT mice the average NMJs’ area stained with BuTx was approximately 330 μm^2^ in males and 250 μm^2^ in females and no change in size was observed between 1 and 3 months of age. In 1-month old DOK7^KI/KI^ mice treated with Amp-101 the NMJs were larger than those of WT littermate controls with the largest being in females treated with the highest dose of 1x10^14^ vg/kg (mean BuTx area of 1242 μm^2^). NMJs in treated model mice increased in size between 1 and 3 months, although this change was not always statistically significant (Fig 4b, one-way ANOVA with Tukey’s multiple comparisons test). We were interested in whether the dose influenced the size of the NMJs, but the only significant difference was in the nerve staining in the female model mice at 1 month of age, with 1x10^14^ vg/kg generating the larger area of 663.4μm^2^ compared with 384μm^2^ for 6x10^13^ vg/kg (Fig 4b, p=0.0367; one-way ANOVA with Tukey’s multiple comparisons test).

In WT NMJs there is a good registration between the pre-and post-synaptic staining. This relative localisation can be quantified as NMJ occupancy by calculating the ratio of pre- and post-synaptic area. For WT mice the NMJ occupancy is approximately 0.7 at 1 month, increasing to around 0.85 by 3 months (Fig 4c). In the treated models at 1 month of age, NMJ occupancy is not significantly different to WT mice. However, at 3 months of age, NMJ occupancy in Amp-101 treated models is significantly lower than for WT littermates (approximately 0.4 for 6x10^13^ vg/kg and 0.45 for 1x10^14^ vg/kg for both genders). This reflects the fact that at 3 months of age the post-synaptic area has increased to a larger extent than the pre-synaptic area, and therefore a lower proportion of the post-synaptic density is covered by the motor neuron terminus. This can be observed in Fig 4a.

### Human DOK7 is expressed in several different muscles

Amp-101 was injected into the peritoneum and so we expected the diaphragm would be readily accessible to the virus, as borne out by the presence of enlarged synapses in this muscle. However, it was important to assess gene expression in other muscles more distal from the injection site. Therefore, the forelimb triceps brachii and hindlimb tibialis anterior from the 3-month old groups were selected for further investigation by western blotting.

DOK7 protein was detected in most of the treated model mice and was virtually undetectable in the WT littermates. However, the levels of expression in treated models varied greatly and this large variation did not seem to correlate with dose. Example blots showing staining of DOK7 are shown in Fig 5a. To semi-quantify DOK7 protein expression, the DOK7 band intensity was normalised to total protein in each lane (visualised by using Revert700 – examples shown in Fig EV2) and then normalised with a positive control loaded onto each gel (Fig 5b). From this data it is clear that Amp-101 injected into the peritoneum is able to transduce muscles that are located in different parts of the body, namely diaphragm, triceps brachii and tibialis anterior, and express DOK7.

**Figure 5.**
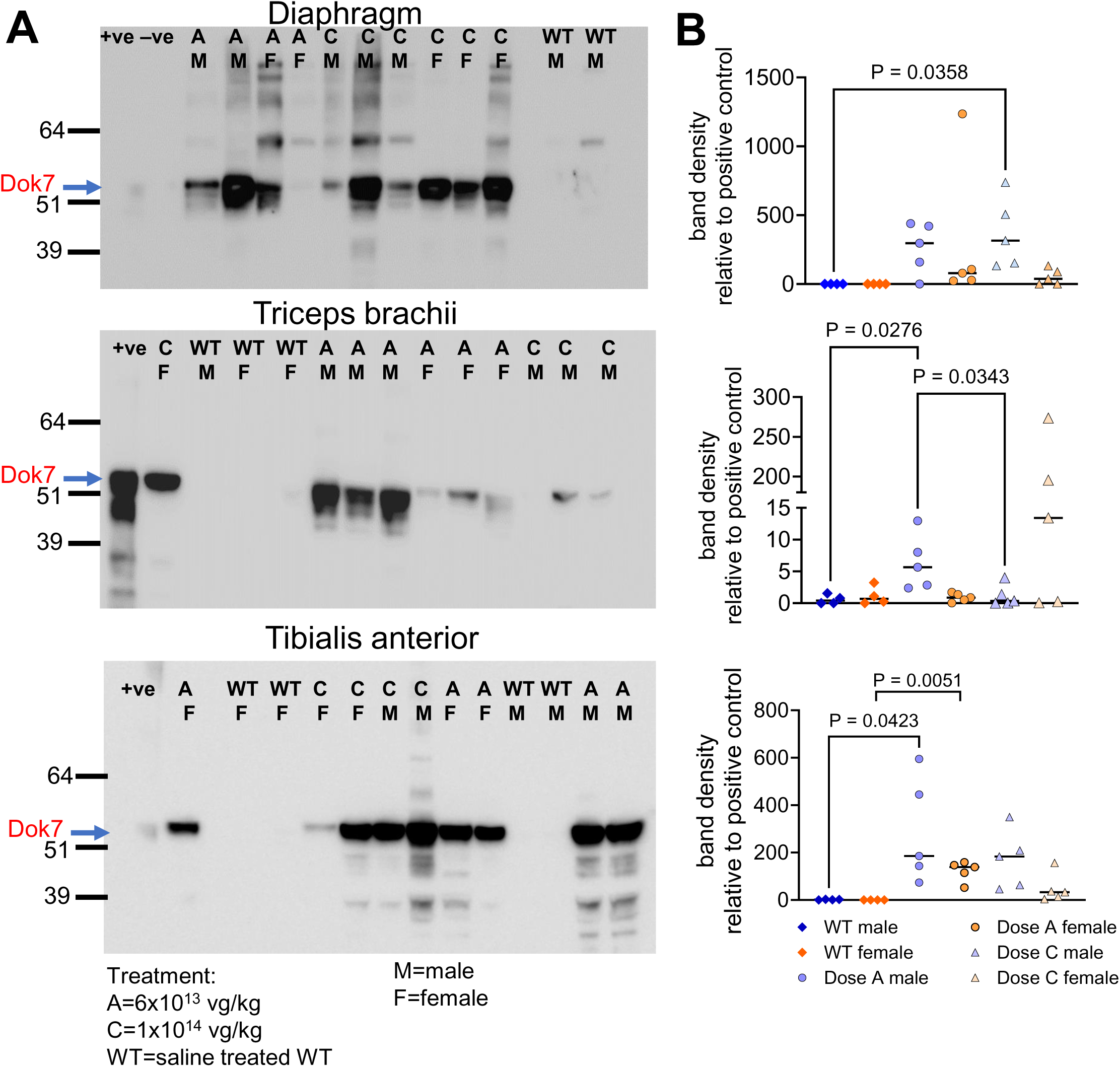
DOK7 is expressed in diaphragm, triceps brachii and TA in Amp-101 treated Dok-7^KI/KI^ model mice. A. Examples of western blots showing DOK7 expression. Robust expression is seen in muscles from many of the treated mice. Endogenous levels of DOK7 in WT mice is very low. B. Densitometry was carried out to semi-quantify DOK7 expression. First the ratio of the band intensity of DOK7 with total protein was calculated, and then this was normalized with a positive control (lysate from HEK292T cells transfected with human DOK7 cDNA). P values were obtained using a one-way ANOVA with Tukey’s multiple comparisons test.

## Discussion

The aim of this project was to test the efficacy of a novel AAV gene therapy, Amp-101, in a severe mouse model of DOK7-CMS. We have demonstrated that when Amp-101 was injected intraperitoneally it could transduce different muscles and generate enlarged synapses. Most remarkably it could rescue this extremely weak and fragile mouse, increasing survival and restoring normal strength.

Recombinant AAV virus is a leading delivery vector of gene therapies and has been widely used in over 250 clinical trials (see (Shen et al, 2022) for meta-analysis). Seven different rAAV gene therapies have reached the market; Roctavian^TM^ (BioMarin) for treatment of haemophilia A and Hemgenix® (uniQure/CSL) for haemophilia B (Dougherty & Dougherty, 2023), Elevidys® (Sarepta Therapeutics) to treat Duchenne muscular dystrophy (Hoy, 2023), Luxturna^TM^ (Spark Therapeutics Inc) for treatment of retinal dystrophy (Askou et al, 2021), Zolgensma® (Novartis Gene Therapies) for spinal muscular atrophy (Waldrop et al, 2020), Glybera® (Uniqure) to treat lipoprotein lipase deficiency (although it is no longer marketed)

(Yla-Herttuala, 2012), and Upstaza^TM^ (PTC Therapeutics) as a treatment for aromatic L-amino acid decarboxylase deficiency (Keam, 2022). Although these successes highlight their therapeutic use, the use of AAV is linked to some adverse effects. rAAV-mediated hepatoxicity is the most common adverse effect and dorsal root ganglia toxicity has been observed after CNS-targeted systemic gene transfer in non-human primates (Hinderer et al, 2018; Hordeaux et al, 2020). Adverse events linked to severe hepatobiliary syndrome and sepsis have caused the death of four children enrolled in the ASPIRO clinical trial (NCT03199469) aiming to show the safety and efficacy of an AAV8 based gene therapy (AT132) for the treatment of X-linked myotubular myopathy (XLMTM) (Wilson & Flotte, 2020). The patients that unfortunately died were older and heavier and thus received among the highest total vg (range: 4.8 × 10^15^ / 7.74 × 10^15^ total vg). Moreover, they had evidence of pre-existing intrahepatic cholestasis which may have played a pathogenic role (D’Amico et al, 2021).

It is therefore preferable to develop rAAV-based therapies with a lower minimum therapeutic dose (MTD). Here, none of the first six animals injected with a dose of 2x10^13^vg/kg after injection survived past two weeks of age, and injections at this dose was stopped on ethical grounds. Successful treatment was observed in 32% and 35% of the model mice injected with 6x10^13^ and 1x10^14^vg/kg respectively. This is lower than the survival rate of 100% described by Arimura et al. when the DOK7^KI/KI^ mouse model was injected with an AAV9 vector in which human DOK7 was under control of the CMV promoter (Arimura *et al*., 2014). It is not clear why 65% of mice reported here failed to thrive, but it could be a combination of several factors. We used an AVVrh74 vector, and although AAVrh74 is able to transduce striated muscle, the AAV9 serotype might have slightly increased early transduction efficiencies (Tabebordbar et al, 2021). Arimura et al. also used a CMV promoter whereas our construct contained a muscle-restricted promoter, which may have different transcription activation kinetics. It is also possible that injection failure might have contributed to mortality, as we were using IP delivery. The most important factor is likely to be the difference in severity of the DOK7^KI/KI^ model in our lab versus in Prof Yamanashi’s lab. Although we obtained the model from Prof Yamanashi, our strain was refreshed before this study by further cross breeding with C57BL/6J mice, so the background strain of the mice used here may be slightly different from that used by Arimura et al. As Oury et al. showed the background strain can modify the severity of the *Dok7* c.1124-1127 dupTGCC knock in genotype, with black six mice having the most severe phenotype (Oury et al, 2021). Unfortunately, our study was initiated prior to this publication, so it was too late to outbreed out strain. Our untreated DOK7^KI/KI^ mice survived fewer than 10 days whereas Arimura et al. reported survival between 13-20 days of age. This meant that there was a shorter window of opportunity for the AAV to transduce the muscle, express DOK7 and remodel the NMJs to become large enough to rescue the phenotype. We injected the pups at 4 days of age to maximise this window, but this still only gave the treatment <5 day to take effect.

Nevertheless, nearly all the animals that did survive past two weeks of age survived for the duration of the study, which is clearly a better outcome than saline injected mice. Moreover, each of the muscle groups we looked at showed effective levels of transduction and, given the survival and strength of the mice, it is likely that all key muscle groups throughout the animals were effectively transduced. It is important to note that this mouse model exhibits a much more severe phenotype than DOK7-CMS patients, and therefore the survival rate for this model does not directly transfer to the clinical context. Having said that, Amp-101 is clearly on a different efficacious level to the current preferred clinical treatment salbutamol, which only extends the lifespan of this mouse model by a maximum of three days (Webster et al, 2020).

Of the mice that survived more than 2 weeks, three males (two injected with 6x10^13^ vg/kg and one injected with 1x10^14^ vg/kg) reached the humane end point for weight loss later in the trial. In addition, one male WT mouse also reached the weight loss humane end point and had to be culled. We could not ascertain a reason for this weight loss, and there was no visible evidence of liver disease, or dental problems that may have affected the ability to eat. It is possible the deaths may have been unrelated to the transgenic status or the therapy, especially since a WT male also lost weight and reached the humane endpoint.

The most important finding from a clinical perspective is that all the mice injected with Amp-101 showed WT levels of strength or better in all the strength tests carried out. Although male mice injected with 1x10^14^ vg/kg were stronger than those injected with 6x10^13^ ^vg/kg^ at the peak of the effect at 6 weeks of age, by the end of the trial both doses gave a similar response to each other, and all the treated model mice were still as strong as WT littermates. For the female treated models there was no difference in strength between the two doses at any age. This data indicates that the MTD for Amp-101 would be around 6x10^13^vg/kg, which is lower than the recommended dose for Zolgensma® (1.1x10^14^vg/kg). It should also be noted that we had to inject Amp-101 intraperitoneally because the 4-day old pups are too small for IV injection, the preferred route of administration in patients. Nonetheless, the relatively low MTD of Amp-101 makes it promising as a potential therapy for DOK7-CMS.

Even though the treated model mice did not show any sign of fatiguability on the inverted screen hang test, EMG analysis indicated that there was a neuromuscular signalling defect in most groups. Decrement indicates a drop-off of muscle fibre recruitment with successive stimuli. As the transduction efficiency of AAVrh74 is not perfect, some fibres may not have been transduced by Amp-101 and might have no NMJs, or some NMJs might be too small to repeatedly generate action potentials. This is supported by the fact that there was a wide variation in NMJ size in all the treated model mice. The fact that the mice are strong suggests that even if this is the case, sufficient muscle fibres are transduced to restore and maintain normal strength. The CMAP amplitude in male mice treated with the highest dose of 1x10^14^ vg/kg did not decrease significantly below that of WT littermates, indicating that this dose can successfully ameliorate decrement.

The AAV genome does not integrates into the cell genome, but remains episomal in the nucleus. Thus, in rapidly dividing cells, the AAV DNA will become diluted and less effective with each cell division. AAV gene therapy is therefore likely to be more effective in tissue that contain slowly or non-dividing cells, such as skeletal muscle. However, the effectiveness may be somewhat negated if the AAV is administered in infancy/childhood (as would be preferred for treatment of CMS), because the muscle will grow and increase in mass, potentially diluting the viral genome. We measured the size of NMJs in treated models at 1 month and 3 months of age, and did not see any evidence of such a dilution effect. In fact, the NMJs increased during this period. Our combined data illustrate that even though the male mice weighed 13 times more, and the females 10 times more, on average by the end of the study compared to when they were injected, Amp-101 continued to induce enough DOK7 expression to maintain NMJ function.

In summary we have demonstrated that Amp-101, a novel rAAV expressing human DOK7, is an effective gene therapy in a mouse model for DOK7-CMS when administered at doses comparable with or lower than current rAAV therapies that are already in the clinic. We also show that the treatment applied at a very young age alleviated the disease phenotype even after the animals grew to adulthood, giving hope that a one-off treatment will bear long lasting effects.

## Methods

### DOK7 model mice

All procedures were performed in accordance with the Animals (Scientific Procedures) Act 1986 and the Home of code of practice and were approved by the University Ethical Review Panel, University of Oxford, UK. Animals were housed at 19–23°C, humidity 45–64% with 12 h light/12 h of dark, in individual ventilated caging (IVC) with aspen chip bedding (Datesand Eco4). They were fed with SDS RM1 food, water was RO chlorinated, and sizzle nest and tubes were included.

Dok-7 knock-in mice homozygous for the common frameshift mutation c.1124_1127dupTGCC were generated by crossing Dok-7^+/KI^ mice with each other. Genotyping was performed by PCR amplification of genomic DNA obtained from tail tips at P3 using primers F1 (ATAGAGGCTGGCTTGGCAGATG) and R1 (TCCTAGCCTAACCATTGTGACTAC) followed by digestion with BamHI which recognises the knocked-in mutation.

### Amp-101

Amp-101 was manufactured by two different companies, Viralgen and Andelyn Biosciences using HEK293 cells in suspensions followed by ultra centrifugation-based purification. All of the mice in the 3-month groups and one from the 1-month group were injected with the rAAV from Viralgen, and the virus from Andelyn Biosciences was used for the rest of the 1-month group. Virus was diluted in sterile 0.9% w/v sodium chloride in low protein binding collection tubes (Life Technologies Ltd) and stored at 4°C. The person making out the dilutions coded the tubes so that the person carrying out the rest of the project and analyses was blinded to the dose. Intraperitoneal injection of 20ul of the Amp-101 into Dok-7^KI/KI^ mice or saline into WT littermates was carried out on P4 using 0.3ml 30g insulin needles (VWR International Ltd). Mice were weighed every day from P3 until P30 after which they were weighed once per week.

### Muscle strength

Muscle strength and fatiguability was assessed using an inverted screen hang test as previously described (Webster et al., 2013; Vanhaesebrouck et al., 2019), a grip strength test (purchased from Bioseb in Vivo Research Instruments) and a rotarod (Omnitech Electronics, Inc). For the inverted screen hang test, mice were placed on a wire mesh which was inverted over a soft padded area and the length of time before the mouse fell was recorded. If the mouse held on for 10 minutes it was removed and allowed 30 seconds rest. This process was carried out three times for each timepoint. The grip test was performed five times at each timepoint and the highest value was used for analysis. The rotarod method used an accelerating profile as follows. The mouse was placed on the rotarod which rotated at 5rpm for 30 seconds, followed by a period of constant acceleration up to 40rpm over a 5-minute period. This was carried out three times for each timepoint, and the highest value was used for analysis.

### Electromyography in mice

Repetitive nerve stimulation was performed as described previously (Webster et al., 2013). Mice were anaesthetised using inhaled Isoflurane (Isoflo anaesthetic 100% w/w, Zoetis inc). Initial induction used 1.5% isoflurane / oxygen, and this was lowered to 1.2% isoflurane / oxygen for maintenance of anaesthesia. Mouse rectal temperature was maintained between 37 °C and 38 °C by using a heat mat underneath the mouse. The sciatic nerve was stimulated at the level of the hip. Compound Muscle Action Potentials (CMAPs) were recorded from gastrocnemius muscles (Dual bio amp/stimulator and Powerlab 4/25, AD Instruments). A train of ten stimuli supramaximal was applied at five different frequencies of 1Hz, 3Hz, 5Hz, 10Hz and 20Hz. Each train was carried out three times. After the EMG procedure was complete, mice were injected sub-cutaneously with 2mg/kg Metacam (Boehringer Ingelheim) and allowed to recover in a heated chamber before returning to the home cage.

The data was analysed using pClamp 9, Molecular Devices.

### Immunofluorescence staining of endplate regions from mouse muscle

Quarter-diaphragms were fixed in 3% formaldehyde (TAAB Laboratory Equipment) in PBS for 1 hour at RT. Tissue was washed 3x with PBS and permeabilised for 1 hour with 0.3% Triton-X in PBS. Tissue was washed again and blocked with 3% bovine serum albumin (BSA) in PBS for 1 hour at RT. Muscle was washed in PBS and incubated overnight at 4°C in PBS containing 3% BSA, rabbit anti-neurofilament heavy chain antibody (Abcam plc, 1:1000) and rabbit anti-synaptophysin Ab-4 (Fisher Scientific, 1:100). Samples were washed extensively with PBS. nAChRs were visualised by incubating muscle with 594-α-BuTx diluted 1:100 for 1 hour at room temperature. Neurofilament and synaptophysin staining was visualised using 488-Alexafluor goat anti rabbit secondary antibody (Fisher Scientific, 1:1000. The tissue was mounted onto microscopy slides using Slowfade Diamond Antifade Mountant (Invitrogen) and z-stacks of neuromuscular junctions were captured using a Zeiss 900 confocal microscope. Maximum projection images were generated and the size of the area staining positive for nAChR or neurofilament and synaptophysin was measured using ImageJ.

### Western blot

#### Tissue preparation

250ul ice-cold extraction buffer (50 mM Tris-HCl pH 7.5, 1% triton X-100 and 500 mM NaCl) supplemented with protease inhibitor cocktail (Sigma, P8340) was added to 2ml tubes containing Precellys Steel beads (2.8 mm ‘reinforced’ 2 ml tubes, pre-filled with steel beads purchased from VWR International Ltd, cat no 432-0142Tubes were kept on ice. Muscle samples were placed into the tubes where they were allowed to thaw on ice. Samples were homogenised using a Precellys 24 lysis homogeniser (Bertin instruments) at 4°C at 6500rpm using a 30-5 seconds program twice. Samples were then rotated for 1 hour at 4°C. The lysate was transferred to clean microcentrifuge tubes which were centrifuged at 13000rpm for 10 mins at 4°C to pellet cell debris. Clarified lysate was transferred to clean microcentrifuge tubes. Samples were diluted typically 1:10, or sometimes 1:20, for protein concentration quantification using a BCA protein assay kit (Fisher Scientific UK Ltd, cat no 10678484).

#### Western blot procedure

20ug protein samples plus NuPAGE LDS sample buffer (Life Technologies Ltd Cat number NP0008) and 10X Bolt™ Sample Reducing Agent (Life Technologies Ltd Cat number B0009) were loaded onto NuPAGE 4 to 12% gradient Bis-Tris, 1.0mm, 15-well mini protein gels (Life Technologies Ltd Cat no NP323BOX). Electrophoresis was carried out at 120V until the dye front reached the bottom of the gel. Protein was transferred to nitrocellulose using NuPAGE transfer buffer (Life Technologies Ltd Cat no NP00061) at 30V for 2 hours.

#### Protein detection antibody staining

Nitrocellulose was dried overnight at room temperature and was rehydrated by incubation in PBS for 5 minutes with gentle rocking and was then rinsed with dH_2_O. It was stained with Revert 700 total protein stain (LI-COR Biosciences UK Ltd cat no 926-11011) for 5 minutes at rt with gentle rocking, followed by washing with wash solution () for 2x30 seconds at rt. Finally, the membrane was rinsed 2x with dH_2_O. Stained membrane was imaged using a Chemidoc imaging system (Bio-Rad Laboratories) using the StarBright B700 settings.

#### Antibody staining

Nitrocellulose was incubated in blocking buffer (PBS containing 3% powdered skimmed milk) for 30 minutes at rt and then incubated with primary mouse anti-DOK7 antibody (Santa Cruz cat no sc-390856) diluted 1:250 in blocking buffer for 1 hour at rt with gentle agitation. Membrane was washed 3x5 minutes with 50mls PBS and then incubated with secondary HRP-conjugated goat anti-mouse antibody (Life Technologies Ltd Cat no 31430) diluted 1:100 in blocking buffer for 1 hour at rt with gentle agitation. Membrane was washed 3x5 minutes with 50mls PBS and HRP activity was detected using ECL according to the manufacturer’s instructions (Merck Life Science UK Limited Cat no GERPN3243). Stained membrane was imaged using a Chemidoc imaging system (Bio-Rad Laboratories) using the Chemiluminescent settings.

Densitometry of the above staining was carried out using Image Lab 6.1.

### Statistics

Graphpad Prism was used for statistical analysis. All error bars shown in figures are standard errors of the mean.

## Acknowledgements

We would like to thank the staff of Biomedical Services for animal care. Thanks also to Yuji Yamanashi for the kind gift of the Dok-7^KI/KI^ mouse model. Research reported in this publication was supported by the National Institute of Neurological Disorders and Stroke (NINDS) of the National Institutes of Health under award number R44NS124351 and funding by Amplo Biotechnology. The content is solely the responsibility of the authors and does not necessarily represent the official views of the National Institutes of Health.

## Author contributions

JC and IK: All of the experimental work and data analysis; writing - reviewing, editing.

JC: Supervision; writing – original draft.

YD: Conceptualisation; supervision; writing - reviewing, editing.

DB: Conceptualisation; writing - reviewing, editing.

PS, CC, AH: Conceptualisation; funding acquisition; writing – reviewing.

CC: Project administration

## The Paper Explained

### Problem

Congenital myasthenic syndromes (CMS) are a rare inherited group of disorders characterised by fatiguable muscle weakness. One of the most commonly occurring CMSs, called DOK7-CMS, is caused by mutations in the *DOK7* gene. Treatment often leads to improvement, but many patients still experience some form of disability, thereby necessitating the development of improved therapies.

### Results

Here we carried out a pre-clinical dose escalation study to assess the efficacy of an innovative adeno-associated virus (AAV) gene therapy called Amp-101 as a potential treatment for this disease. The DOK7-CMS mouse model we used is very weak and only survives a few days if untreated. Following a single treatment with Amp-101 at 4 days of age, around 35% of the mice became as strong as wild type mice and survived until the end of the trial.

### Impact

Amp-101 is an effective gene therapy in our mouse model for DOK7-CMS and is a promising candidate to move forward to clinical trials. The results give hope that a single treatment administered in childhood will bear long-lasting effects.

## References

1. (2020) High-dose AAV gene therapy deaths. *Nat Biotechnol* 38: 910

2. Arimura S, Okada T, Tezuka T, Chiyo T, Kasahara Y, Yoshimura T, Motomura M, Yoshida N, Beeson D, Takeda S, Yamanashi Y (2014) Neuromuscular disease. DOK7 gene therapy benefits mouse models of diseases characterized by defects in the neuromuscular junction. Science 345: 1505–1508

3. Armstrong A, 2021. Pfizer reports patient death in early-stage Duchenne gene therapy trial, halts enrollment.

4. Askou AL, Jakobsen TS, Corydon TJ (2021) Retinal gene therapy: an eye-opener of the 21st century. Gene Ther 28: 209–216

5. Beeson D, Higuchi O, Palace J, Cossins J, Spearman H, Maxwell S, Newsom-Davis J, Burke G, Fawcett P, Motomura M et al (2006) Dok-7 mutations underlie a neuromuscular junction synaptopathy. Science 313: 1975–1978

6. Burke G, Hiscock A, Klein A, Niks EH, Main M, Manzur AY, Ng J, de Vile C, Muntoni F, Beeson D, Robb S (2013) Salbutamol benefits children with congenital myasthenic syndrome due to DOK7 mutations. Neuromuscul Disord 23: 170–175

7. Clausen L, Cossins J, Beeson D (2018) Beta-2 Adrenergic Receptor Agonists Enhance AChR Clustering in C2C12 Myotubes: Implications for Therapy of Myasthenic Disorders. J Neuromuscul Dis 5: 231–240

8. D’Amico A, Longo A, Fattori F, Tosi M, Bosco L, Chiarini Testa MB, Paglietti MG, Cherchi C, Carlesi A, Mizzoni I, Bertini E (2021) Hepatobiliary disease in XLMTM: a common comorbidity with potential impact on treatment strategies. Orphanet journal of rare diseases 16: 425

9. Dougherty JA, Dougherty KM (2023) Valoctocogene Roxaparvovec and Etranacogene Dezaparavovec: Novel Gene Therapies for Hemophilia A and B. Ann Pharmacother: 10600280231202247

10. Eguchi T, Tezuka T, Fukudome T, Watanabe Y, Sagara H, Yamanashi Y (2020) Overexpression of Dok-7 in skeletal muscle enhances neuromuscular transmission with structural alterations of neuromuscular junctions: Implications in robustness of neuromuscular transmission. Biochem Biophys Res Commun 523: 214–219

11. Hinderer C, Katz N, Buza EL, Dyer C, Goode T, Bell P, Richman LK, Wilson JM (2018) Severe Toxicity in Nonhuman Primates and Piglets Following High-Dose Intravenous Administration of an Adeno-Associated Virus Vector Expressing Human SMN. Hum Gene Ther 29: 285–298

12. Hordeaux J, Buza EL, Jeffrey B, Song C, Jahan T, Yuan Y, Zhu Y, Bell P, Li M, Chichester JA et al (2020) MicroRNA-mediated inhibition of transgene expression reduces dorsal root ganglion toxicity by AAV vectors in primates. Sci Transl Med 12

13. Hoy SM (2023) Delandistrogene Moxeparvovec: First Approval. Drugs 83: 1323–1329

14. Keam SJ (2022) Eladocagene Exuparvovec: First Approval. Drugs 82: 1427–1432

15. Huang YT, Crick HR, Chaytow H, van der Hoorn D, Alhindi A, Jones RA, Hector RD, Cobb SR, Gillingwater TH (2023) Long-term muscle-specific overexpression of DOK7 in mice using AAV9-tMCK-DOK7. Mol Ther Nucleic Acids 33: 617–628

16. Inoue A, Setoguchi K, Matsubara Y, Okada K, Sato N, Iwakura Y, Higuchi O, Yamanashi Y (2009) Dok-7 activates the muscle receptor kinase MuSK and shapes synapse formation. Sci Signal 2: ra7

17. Kaifer KA, Villalon E, Smith CE, Simon ME, Marquez J, Hopkins AE, Morcos TI, Lorson CL (2020) AAV9-DOK7 gene therapy reduces disease severity in Smn(2B/-) SMA model mice. Biochem Biophys Res Commun 530: 107–114

18. Lashley D, Palace J, Jayawant S, Robb S, Beeson D (2010) Ephedrine treatment in congenital myasthenic syndrome due to mutations in DOK7. Neurology 74: 1517–1523

19. Lee M, Beeson D, Palace J (2018) Therapeutic strategies for congenital myasthenic syndromes. Ann N Y Acad Sci 1412: 129–136s

20. Liewluck T, Selcen D, Engel AG (2011) Beneficial effects of albuterol in congenital endplate acetylcholinesterase deficiency and Dok-7 myasthenia. Muscle Nerve 44: 789–794

21. Lorenzoni PJ, Scola RH, Kay CS, Filla L, Miranda AP, Pinheiro JM, Chaouch A, Lochmuller H, Werneck LC (2013) Salbutamol therapy in congenital myasthenic syndrome due to DOK7 mutation. Journal of the neurological sciences 331: 155–157

22. Mahjneh I, Lochmuller H, Muntoni F, Abicht A (2013) DOK7 limb-girdle myasthenic syndrome mimicking congenital muscular dystrophy. Neuromuscul Disord 23: 36–42

23. Mendell JR, Sahenk Z, Lehman K, Nease C, Lowes LP, Miller NF, Iammarino MA, Alfano LN, Nicholl A, Al-Zaidy S et al (2020) Assessment of Systemic Delivery of rAAVrh74.MHCK7.micro-dystrophin in Children With Duchenne Muscular Dystrophy: A Nonrandomized Controlled Trial. JAMA Neurol 77: 1122–1131

24. Miyoshi S, Tezuka T, Arimura S, Tomono T, Okada T, Yamanashi Y (2017) DOK7 gene therapy enhances motor activity and life span in ALS model mice. EMBO Mol Med 9: 880–889

25. Ohno K, Ohkawara B, Shen XM, Selcen D, Engel AG (2023) Clinical and Pathologic Features of Congenital Myasthenic Syndromes Caused by 35 Genes-A Comprehensive Review. Int J Mol Sci 24: 3730

26. Okada K, Inoue A, Okada M, Murata Y, Kakuta S, Jigami T, Kubo S, Shiraishi H, Eguchi K, Motomura M et al (2006) The muscle protein Dok-7 is essential for neuromuscular synaptogenesis. Science 312: 1802–1805

27. Oury J, Zhang W, Leloup N, Koide A, Corrado AD, Ketavarapu G, Hattori T, Koide S, Burden SJ (2021) Mechanism of disease and therapeutic rescue of Dok7 congenital myasthenia. Nature 595: 404–408

28. Palace J, Lashley D, Newsom-Davis J, Cossins J, Maxwell S, Kennett R, Jayawant S, Yamanashi Y, Beeson D (2007) Clinical features of the DOK7 neuromuscular junction synaptopathy. Brain 130: 1507–1515

29. Philippidis A (2022) Novartis Confirms Deaths of Two Patients Treated with Gene Therapy Zolgensma. Hum Gen Ther 33: 842–844

30. Pozsgai ER, Griffin DA, Heller KN, Mendell JR, Rodino-Klapac LR (2017) Systemic AAV-Mediated beta-Sarcoglycan Delivery Targeting Cardiac and Skeletal Muscle Ameliorates Histological and Functional Deficits in LGMD2E Mice. Molecular therapy : the journal of the American Society of Gene Therapy 25: 855–869

31. Rodriguez Cruz PM, Palace J, Beeson D (2018) The Neuromuscular Junction and Wide Heterogeneity of Congenital Myasthenic Syndromes. Int J Mol Sci 19: 1677

32. Schara U, Barisic N, Deschauer M, Lindberg C, Straub V, Strigl-Pill N, Wendt M, Abicht A, Muller JS, Lochmuller H (2009) Ephedrine therapy in eight patients with congenital myasthenic syndrome due to DOK7 mutations. Neuromuscul Disord 19: 828–832

33. Shen W, Liu S, Ou L (2022) Corrigendum: rAAV immunogenicity, toxicity, and durability in 255 clinical trials: A meta-analysis. Front Immunol 13: 1104646

34. Slater CR, Fawcett PR, Walls TJ, Lyons PR, Bailey SJ, Beeson D, Young C, Gardner-Medwin D (2006) Pre- and post-synaptic abnormalities associated with impaired neuromuscular transmission in a group of patients with ‘limb-girdle myasthenia’. Brain 129: 2061–2076

35. Sondergaard PC, Griffin DA, Pozsgai ER, Johnson RW, Grose WE, Heller KN, Shontz KM, Montgomery CL, Liu J, Clark KR et al (2015) AAV.Dysferlin Overlap Vectors Restore Function in Dysferlinopathy Animal Models. Ann Clin Transl Neurol 2: 256–270

36. Tabebordbar M, Lagerborg KA, Stanton A, King EM, Ye S, Tellez L, Krunnfusz A, Tavakoli S, Widrick JJ, Messemer KA et al (2021) Directed evolution of a family of AAV capsid variants enabling potent muscle-directed gene delivery across species. Cell 184: 4919–4938 e4922

37. Ueta R, Sugita S, Minegishi Y, Shimotoyodome A, Ota N, Ogiso N, Eguchi T, Yamanashi Y (2020) DOK7 Gene Therapy Enhances Neuromuscular Junction Innervation and Motor Function in Aged Mice. iScience 23: 101385

38. Vanhaesebrouck AE, Webster R, Maxwell S, Rodriguez Cruz PM, Cossins J, Wickens J, Liu WW, Cetin H, Cheung J, Ramjattan H et al (2019) beta2-Adrenergic receptor agonists ameliorate the adverse effect of long-term pyridostigmine on neuromuscular junction structure. Brain 142: 3713–3727

39. Waldrop MA, Karingada C, Storey MA, Powers B, Iammarino MA, Miller NF, Alfano LN, Noritz G, Rossman I, Ginsberg M et al (2020) Gene Therapy for Spinal Muscular Atrophy: Safety and Early Outcomes. Pediatrics 146

40. Webster RG, Vanhaesebrouck AE, Maxwell SE, Cossins JA, Liu W, Ueta R, Yamanashi Y, Beeson DMW (2020) Effect of salbutamol on neuromuscular junction function and structure in a mouse model of DOK7 congenital myasthenia. Hum Mol Genet 29: 2325–2336

41. Wilson JM, Flotte TR (2020) Moving Forward After Two Deaths in a Gene Therapy Trial of Myotubular Myopathy. Hum Gene Ther 31: 695–696

42. Yla-Herttuala S (2012) Endgame: glybera finally recommended for approval as the first gene therapy drug in the European union. Molecular therapy : the journal of the American Society of Gene Therapy 20: 1831–1832

43. Zygmunt DA, Crowe KE, Flanigan KM, Martin PT (2017) Comparison of Serum rAAV Serotype-Specific Antibodies in Patients with Duchenne Muscular Dystrophy, Becker Muscular Dystrophy, Inclusion Body Myositis, or GNE Myopathy. Hum Gene Ther 28: 737–746

